# Structural rearrangements and selection promote phenotypic evolution in *Anolis* lizards

**DOI:** 10.1101/2024.12.12.628123

**Authors:** Raúl Araya-Donoso, Sarah M. Baty, Jaime E. Johnson, Eris Lasku, Jody M. Taft, Rebecca E. Fisher, Jonathan Losos, Greer A. Dolby, Kenro Kusumi, Anthony J. Geneva

**Affiliations:** School of Life Sciences, Arizona State University, Tempe, AZ 85281 USA; School of Animal, Plant and Environmental Sciences, University of the Witwatersrand, Johannesburg, South Africa; Department of Biology, Washington University, Saint Louis, MO, USA; Department of Biology, University of Alabama at Birmingham, AB, USA; Department of Biology & Center for Computational and Integrative Biology, Rutgers University–Camden, Camden, NJ 08103 USA

## Abstract

The genomic characteristics of adaptively radiated groups could contribute to their high species number and ecological disparity, by increasing their evolutionary potential. Here, we explored the genomic features of *Anolis* lizards, focusing on three species with unique phenotypes: *A. auratus*, one of the species with the longest tail; *A. frenatus*, one of the largest species; and *A. carolinensis*, one of the species that inhabits the coldest environments. We assembled and annotated two new chromosome-level reference genomes for *A. auratus* and *A. frenatus*, and compared them with the available genomes of *A. carolinensis* and *A. sagrei*. We evaluated the presence of structural rearrangements, quantified the density of repeat elements, and identified signatures of positive selection in coding and regulatory regions. We detected substantial rearrangements in scaffolds 1, 2 and 3 of *A. frenatus* different from the other species, in which the rearrangement breakpoints corresponded to hotspots of developmental genes. Further, we detected an accumulation of repeats around key developmental genes in anoles and phrynosomatid outgroups. Finally, we detected signatures of positive selection on coding sequences and regulatory regions of genes relevant to development and physiology that could affect the unique phenotypes of the analyzed species. Our results suggest that anoles have genomic features associated with genes that affect organismal morphology and physiology. This could provide a genomic substrate that promoted phenotypic disparity in anoles, and contributed to their ability to adaptively radiate.

**Author Summary:** Adaptive radiations are often characterized by high species richness and phenotypic differentiation. Besides the ecological context, the genetic features of organisms could also contribute to their ability to diversify. *Anolis* lizards are an adaptively radiated group that shows high phenotypic disparity in morphology and physiology. In this study, we explored the genome of four species within the *Anolis* radiation with distinctive phenotypes. We generated a high-quality chromosome-level reference genome for *A. auratus* and *A. frenatus*, and compared them with *A. carolinensis* and *A. sagrei*. We detected major structural rearrangements in *A. frenatus*, a high density of repeat elements around key developmental genes, and signatures of natural selection associated with genes functionally relevant for the analyzed species. Hence, the genomic characteristics of anoles were associated with their unique phenotypic diversity. We highlight the potential relevance of genomic features to influence the ability of groups of organisms to radiate adaptively.

## Introduction

Adaptively radiated groups of organisms are natural experiments in which the relative roles of ecological and genomic features on speciation and phenotypic differentiation can be assessed (1–3). In general, ecological factors and the emergence of ecological opportunity are known to play an important role in determining the ability of a group of organisms to radiate adaptively (4,5). On the other hand, genetic mechanisms could also influence the ability of organisms within radiations to diversify and generate extensive phenotypic variation because clades with greater evolutionary potential could be more likely to radiate adaptively (1,6). Multiple genetic mechanisms could contribute to increased genetic and phenotypic diversity such as chromosome-level structural rearrangements, small-scale structural variation, the dynamics of transposable elements, mutation rates, recombination rates, and the genomic landscape of selection on regulatory elements and/or coding regions (6–10).

The relevance of the genomic substrate for highly speciose or adaptively radiated groups of organisms has been discussed before. For example, African lake cichlids show ancient genetic polymorphisms, structural rearrangements, high divergence in regulatory sequences, insertion of transposable elements within regulatory elements, and novel miRNAs (6,8,11). Darwin’s finches also exhibit evidence of ancient polymorphisms, and selection on large-effect loci associated with beak morphology located in genomic islands of low recombination (9,12). *Heliconius* butterflies present increased genomic variation by hybridization and/or introgression processes, high variability in regulatory regions, genome expansion events caused by an increase in repeat elements, and structural rearrangements (13–16). Therefore, groups that radiate could have more labile genomes that allow for greater phenotypic diversification. A current challenge is to determine the relative importance of each of these genomic factors, and whether different radiations present similar genomic features that aided diversification or whether different radiations have occurred through different genomic mechanisms.

*Anolis* lizards are an ideal group to assess the relevance of genetic mechanisms for generating and promoting phenotypic diversity. This genus is described as an adaptive radiation with ∼400 species distributed in the tropical Americas (17,18). *Anolis* are considered a model system for evolutionary biology studies because they present extensive phenotypic variation across multiple niche axes. A remarkable characteristic of *Anolis* evolution is the repeated occurrence of intra-island radiation and morphological differentiation associated with microhabitat use patterns (19–21). Besides morphology, anoles have diversified in behavior, physiology, and sexual dimorphism (22–24). In this context, anoles present a wide range of phenotypic variation compared to other taxa and this diversity may be promoted by ecological and genetic mechanisms.

Within the *Anolis* radiation, some species have distinctive phenotypes that could be adaptive to their niches (Fig. 1A). For example, *A. frenatus* is one of the largest anole species (Fig. 1B), which may reduce its predation risk and enable a wider dietary breadth, potentially including other anole lizards as prey (25); *A. auratus* inhabits grasslands and perches on narrow branches and features an extremely long tail (Fig. 1C) that may provide better balance when walking and jumping along narrow surfaces (26,27); and *A. carolinensis* is one of the species with highest cold tolerance (Fig. 1D), enabling its colonization towards higher latitudes and survival during cold seasons (28). Different types of genomic variation, particularly within coding regions, may control such traits. For instance, longer tails could be produced by modifications of the number and/or size of the caudal vertebrae, controlled by molecular pathways involved in the axial skeleton development (29–31). A larger body size could be controlled by insulin growth factor or growth hormone pathways (32–34), while cold adaptation could be related to genes regulating oxygen consumption and/or blood circulation (28,35).

**Figure 1.**
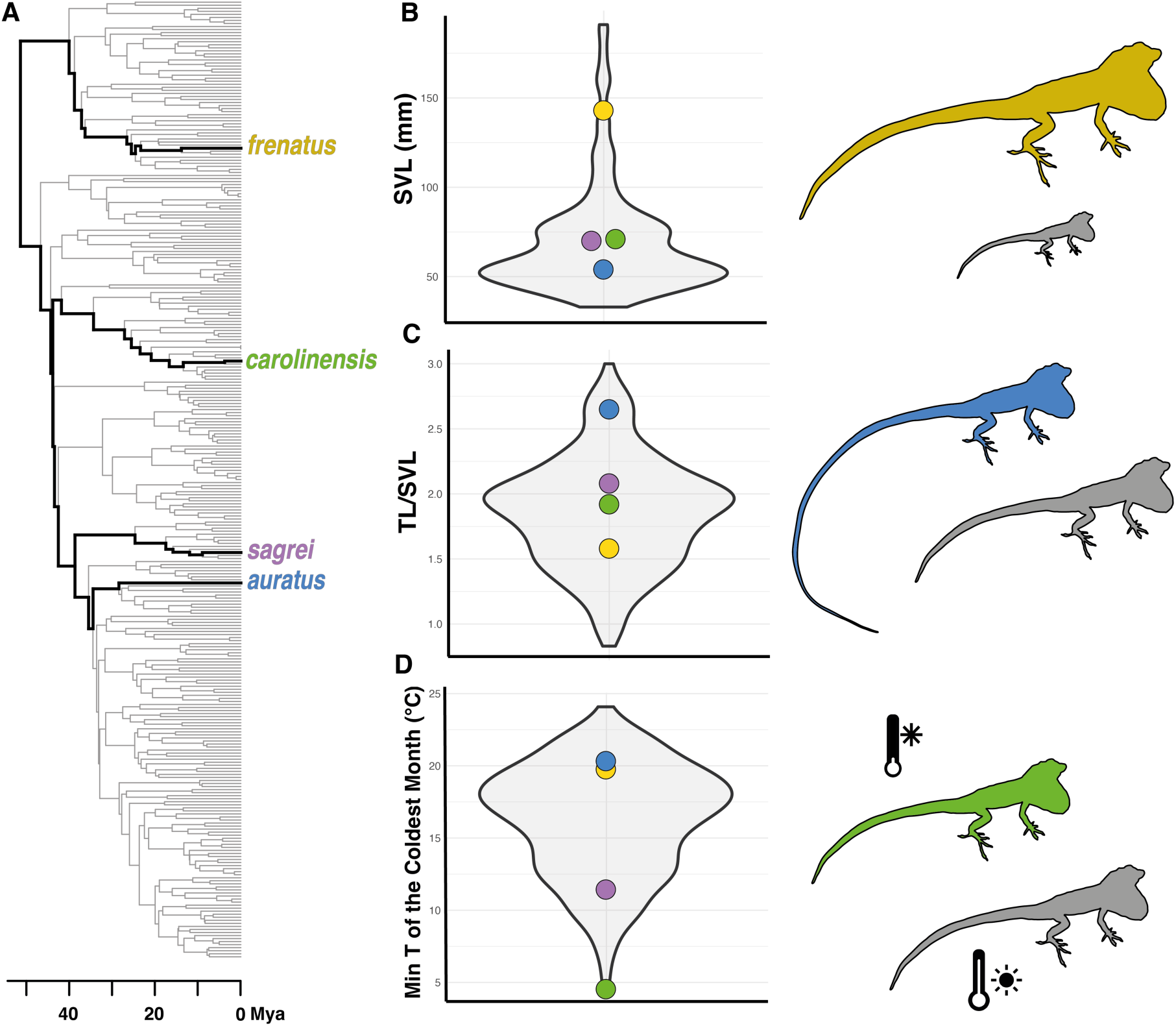
*Anolis* phylogenetic relationships (**A**) and genus-wide phenotypic variation in snout-vent length (SVL; **B**), tail length (TL; **C**), and thermal climatic niche (**D**), highlighting the species included in this study (Phylogenetic and morphological data from Poe et al., (2017); temperature data obtained from WorldClim 2 (37) for all species).

Some genomic features could be associated with the high ecological disparity described in these *Anolis* species. Tollis et al. (2018) compared short-read genome assemblies of five species (including *A. carolinensis*, *A. auratus* and *A. frenatus*), and detected high mutation rates in anoles compared to other vertebrates, and signatures of natural selection on genes associated with limb and brain development and hormonal regulation. In some Cuban anole species, an accumulation of gene duplications has been reported (39), and genomic regions undergoing accelerated evolution have been identified in association with thermal biology (40). Furthermore, *Anolis* genetic diversity could have been fueled by ancient hybridization and introgression processes (41,42). Chromosomal rearrangements could also be relevant because multiple events of chromosome gains and losses have been described within *Anolis* (43,44), and chromosome fission and fusions have been proposed to determine the evolution of the *Anolis* X chromosome (45,46). Finally, the dynamics of repeat elements could be relevant because transposable elements can impact the genome by modifying gene regulation patterns, causing mutations, or promoting genome rearrangements (7). A high density of transposable elements within the *hoxB and hoxC* gene clusters, key regulators of morphological development, has been reported in *Anolis* (47,48). Nonetheless, genome-wide patterns associated with repeat density and chromosome-level structural rearrangements remain to be explored with a genomic approach, because the analysis of those features requires highly contiguous genome assemblies.

Here, we explored the genomic features of species with disparate phenotypes within the adaptively radiated *Anolis* group. We generated chromosome-level reference genomes for two *Anolis* species and found evidence for major structural rearrangements, described a unique pattern of repeat density through the genome, and identified genes under positive selection that were associated with the unique traits from four species representing divergent phenotypes. These genomic features could fuel genetic diversity and hence, promote the high diversification and phenotypic disparity found within *Anolis*.

## Results

### Chromosome level genome assemblies and annotation for A. auratus and A. frenatus

We generated Hi-C chromosome-level genome assemblies for two *Anolis* species (Table 1). Both type specimens were adult females from Panama (Table S1). The resulting assemblies were highly contiguous (N50: *A. auratus* 281.8 Mbp and *A. frenatus* 342.7 Mbp) and complete (BUSCO eukaryotic completeness of 93.07% for *A. auratus* and 86.14 % for *A. frenatus*). Both species show a similar pattern of repetitive element composition (Fig. S1), which corresponds to roughly 50% of the genome. However, *A. auratus* shows a recent accumulation of LINEs. We generated genome annotations for both species via the MAKER pipeline (49) using a combination of new RNA sequencing data, and the proteomes of previously sequenced species (see Methods for details). For *A. auratus* we identified 19,879 genes with an average length of 19,877 bp (Table S2), and 88.2% of all eukaryote BUSCO genes present in the annotation (either complete or fragmented), whereas for *A. frenatus* 19,643 genes were identified with an average length of 18,033 bp (Table S2) and 76.1% eukaryotic BUSCO genes present. For subsequent analyses, our newly annotated genomes were compared against the chromosome-level reference genomes of *A. carolinensis* (AnoCar2.0, (50); and DNAzoo Hi-C Assembly, (51,52)) and *A. sagrei* (AnoSag2.1, (45), along with the phrynosomatid lizards *Urosaurus nigricaudus* (53) and *Phrynosoma platyrhinos* (54).

**Table 1.** Genome assembly and annotation statistics for the four analyzed *Anolis* species.

| Species | *A. auratus* | *A. frenatus* | *A. carolinensis* | *A. sagrei* |
| --- | --- | --- | --- | --- |
| Genome version | RUC_Aaur_2 | RUC_Afre_2 | AnoCar2.0 | AnoSag2.1 |
| Assembly length (Gbp) | 1.77 | 1.85 | 1.79 | 1.66 |
| N50 (Mbp) | 281.8 | 342.7 | 150.6 | 253.6 |
| L50 (n°) | 3 | 3 | 5 | 4 |
| Eukaryote BUSCO Assembly (%) | C + F: 93.07 | C + F: 86.14 | C + F: 94.5% | C + F: 100% |
| % Repeats | 48.53 | 51.27 | 33 | 46.3 |
| N° genes | 19,879 | 19,643 | 21,555 | 20,033 |
| Average gene length (bp) | 19,877 | 18,033 | 32,969 | 45,059 |
| Eukaryote BUSCO Annotation (%) | C + F: 88.2 | C + F: 76.1 | C + F: 94.5% | C + F: 99.7% |
| Reference | This study | This study | Alföldi et al. 2011 | Geneva et al. 2022 |

### Chromosome-level structural rearrangements

We performed *in silico* chromosome painting to assess the synteny conservation among our four *Anolis* species and *U. nigricaudus* and *P. platyrhinos*, using *A. sagrei* as a reference. Overall, there is high synteny conservation for the main scaffolds or macrochromosomes among those species (Fig. 2A). Interestingly, scaffolds 1, 2, and 3 contain substantial structural rearrangements that are unique to *A. frenatus* (Fig. 2A, Fig. 2B). The Hi-C data for *A. frenatus* shows higher contact density within scaffolds and very little interaction between scaffolds 1, 2 and 3 (Fig. 2C, Fig. S2). This observation suggests that the observed rearrangements are not a sequencing or scaffolding artifact, but rather supports genuine structural differences in this species relative to other Iguanian taxa. To establish a link between the structural rearrangements and their potential functional implications, we identified the genes located within 1 Mbp to the rearrangement breakpoints in scaffolds 1, 2 and 3 between *A. sagrei* and *A. frenatus* (Table S3). We conducted an enrichment analysis on the list of genes co-located to the breakpoints with g:Profiler (55) which showed significant enrichment of biological processes such as "cellular differentiation", "developmental process" and "pigment granule transport" (Table S4). Further, we quantified the density of genes associated with developmental GO terms along scaffolds 1, 2 and 3 of *A. frenatus*, and we detected that the chromosomal breaks were located in hotspots of genes with developmental functions (Fig. 2D). Among the genes contiguous to the rearrangement breakpoints (Table S3) we identified *axin2*, a regulator of the Wnt/β-catenin and TGF-β pathways that determines chondrocyte maturation and axial skeletal development (56); *bmp2*, a growth factor determinant for bone development through the BMP-Smad pathway (57); *ddit3*, transcription factor that influences myogenesis by regulating the GH-IGF1 pathway (58); and *twist2*, a transcription factor relevant for bone formation and myogenesis (59).

**Figure 2.**
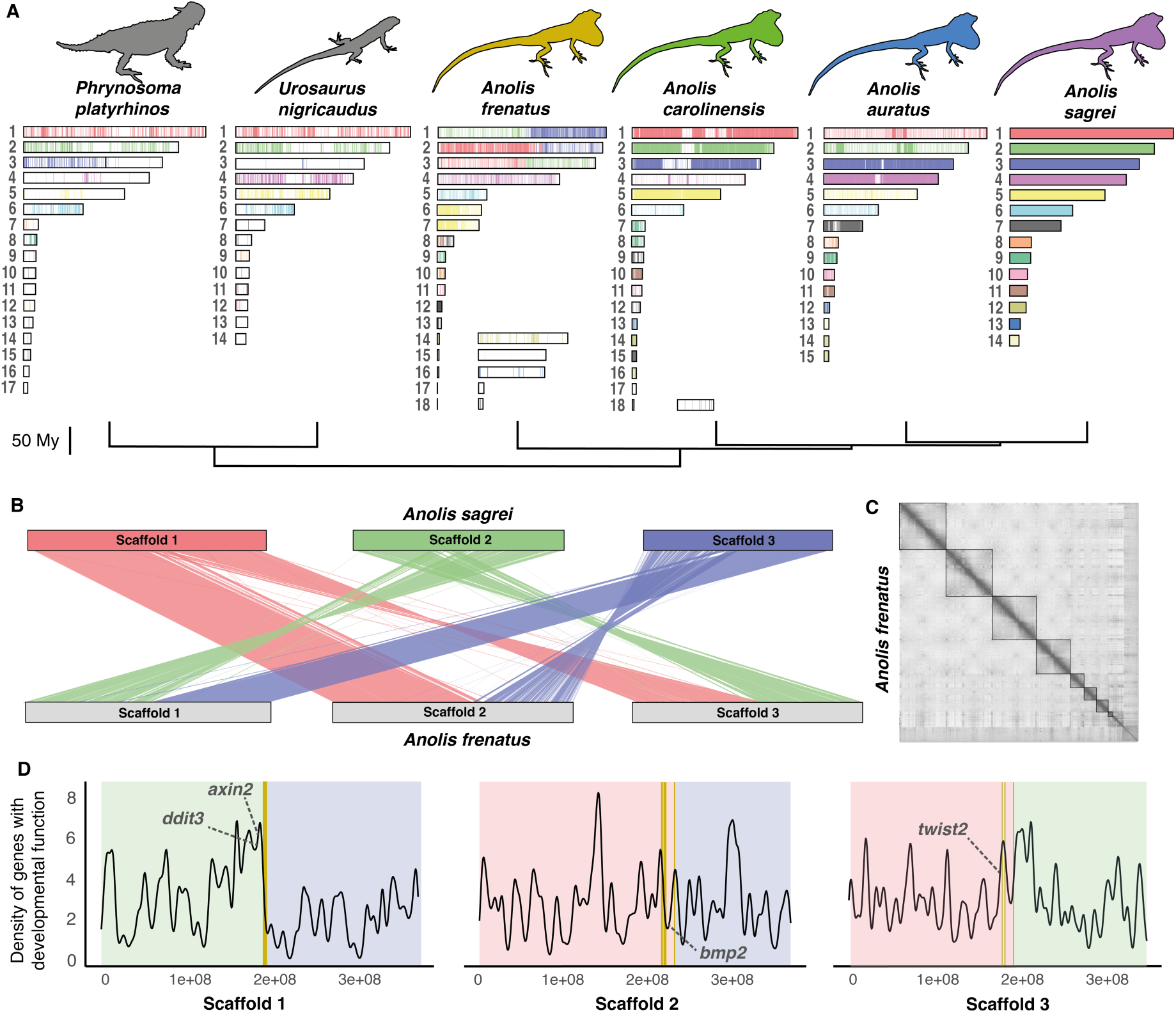
Chromosome-level structural variation across *Anolis*. **A.** Synteny between *A. sagrei* and other anole (*A. auratus*, *A. carolinensis*, *A. frenatus*) and lizard (*U. nigricaudus*, *P. platyrhinos*) species for the largest scaffolds representing the chromosomes of each species. **B.** Synteny between scaffolds 1, 2, and 3 of *A. sagrei* and *A. frenatus* showing substantial rearrangements. **C.** Hi-C density contact matrix for *A. frenatus*. **D.** Density of genes associated with developmental GO terms along scaffolds 1, 2 and 3 in *A. frenatus*. Background colors indicate the homology to *A. sagrei* scaffolds for different chromosomal regions and vertical lines indicate the chromosomal breakpoints. Rearrangement breakpoints are within hotspots of developmental genes.

Scaffold 7 in *A. sagrei* has previously been hypothesized to be the X chromosome and the result of a series of autosomal fusions (45,46,60). *A. auratus* and *A. sagrei* belong to the *Norops* clade of *Anolis* (36) and we found a high degree of synteny conservation between scaffold 7 of these two species, whereas in the species outside of the *Norops* clade it corresponded to a series of smaller scaffolds (Fig. 2A, Fig. S3). To further explore scaffold 7 evolution within anoles we compared this chromosome against another recently published *Norops* clade high-quality genome, *A. apletophallus* (61), which also revealed high synteny conservation with both *A. sagrei* and *A. auratus* (Fig. S3).

### Repeat density is associated with key developmental genes in Anolis and other pleurodonts

We estimated the density of repeats in 500 kb windows throughout the first 6 scaffolds of *A. frenatus*, *A. auratus*, *A. sagrei*, *U. nigricaudus* and *P. platyrhinos* based on the repeat element annotation of their genomes. We selected the densest repeat regions corresponding to the top 5% of repeat density and identified the genes present in those regions from our annotations (Table S5). Within repeat-rich regions in all the analyzed species we detected some developmental genes (Fig. 3) such as the *hoxB*, *hoxC*, and *hoxD* gene clusters, key determinants of the vertebrate body plan (30); *notch4*, a member of the NOTCH receptors family that are crucial for development (62); and *fgf11*, member of the fibroblast growth factor (FGF) family which are involved in development and morphogenesis (63). An enrichment analysis was conducted on the lists of genes located within these high repeat-density regions for each species with g:Profiler. In general, genes associated with regulatory and developmental biological processes were overrepresented in the high repeat-density regions for all species (Table S6; Fig. S4).

**Figure 3.**
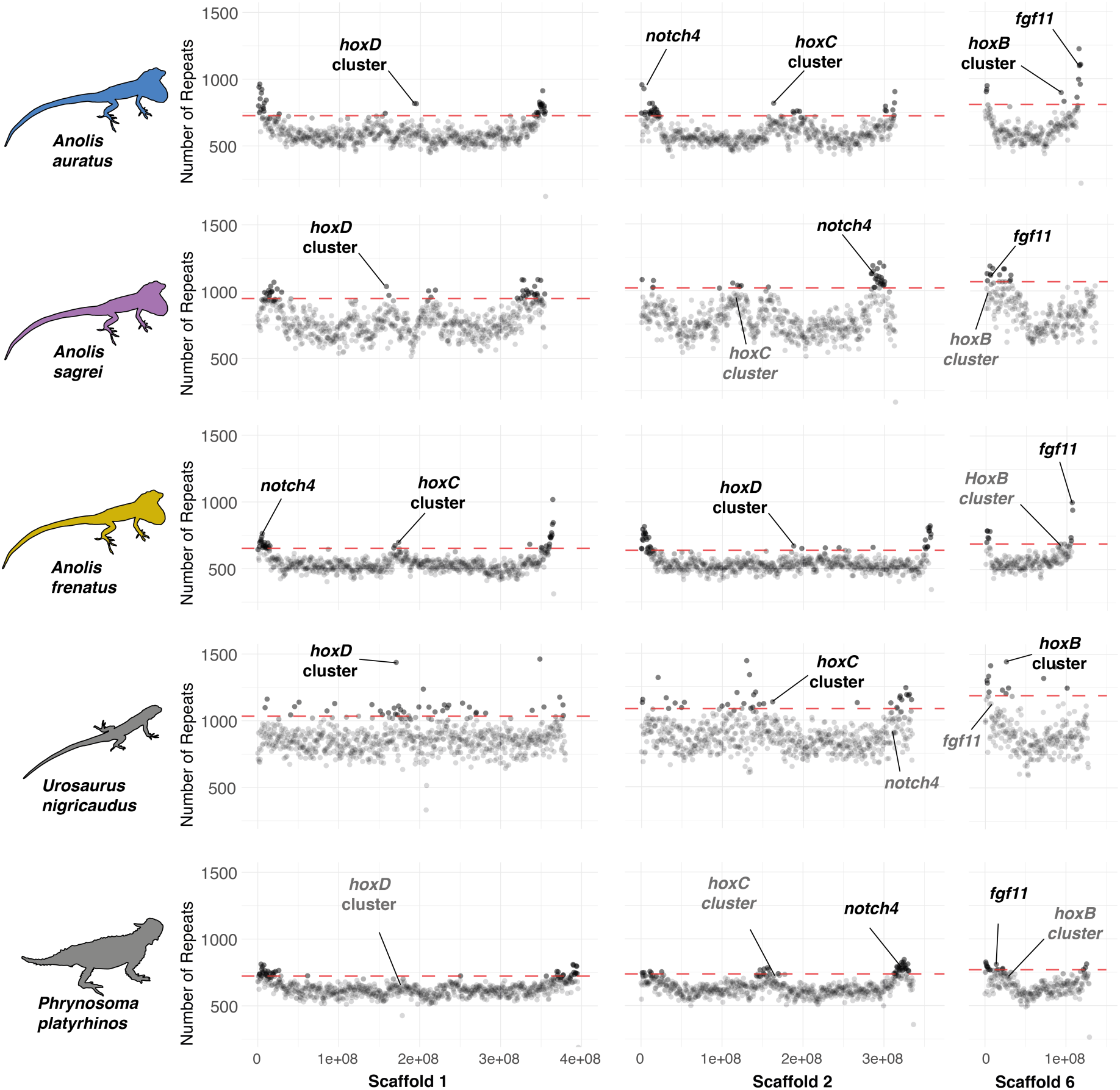
Number of repeat elements in 500 kb windows throughout scaffolds 1, 2 and 6 in the analyzed pleurodont species. A higher density of repeats is found close to key developmental genes in the four *Anolis* and the outgroups.

### Genes under natural selection and divergence in regulatory regions

We identified genes under positive selection among the four *Anolis* species by calculating the pairwise ratio between non-synonymous to synonymous substitutions (dN/dS) between each species pair (Table S7). We retained genes with dN/dS > 1 overlapping in at least 2 out of 3 comparisons for each species (Fig. S5). For *A. frenatus* 16 genes overlapped including *mtpn*, a muscle growth factor that shows similar effects to *igf1* (64,65); and *pdzk1ip1* that regulates and inhibits transforming growth factor (TGF-β) and bone morphogenic protein (BMP) signaling (66). For *A. auratus* 12 genes overlapped including *ramp2* which regulates angiogenesis, cardiovascular development, and influences bone formation (67,68); and *dcdc1*, associated with bone mineral density (69) and bone degradation in humans (70). In *A. carolinensis* we detected 6 overlapping genes including *lep*, a gene relevant to lipid metabolism and energetic balance, and that has thermogenic effects on skeletal muscle (71–73); *clps*, involved in lipid digestion (74); and *stard6*, associated with the intracellular transport of sterol and other lipids (75). *Anolis sagrei* presented 8 overlapping genes including *ppdpf1*, associated with cell proliferation in multiple types of cancer (76); and *s100a1*, that can regulate cell growth and proliferation (77). A gene enrichment analysis was run with g:Profiler for each species (Table S8).

Among the overrepresented GO terms for *A. carolinensis* we detected “lipid catabolic process” and “digestion”, for *A. auratus* “positive regulation of developmental processes”, for *A. frenatus* “regulation of muscle organ development”, and for *A. sagrei* “regulation of polarized epithelial cell differentiation”.

To identify diverged regulatory regions, we identified genes with the top 1% of divergence in their putative promoter regions (1,000 bp upstream of the transcription start site, (78) for each species pair (Table S9). We retained genes that overlapped in at least 2 out of 3 species comparisons. Within the overlapping genes identified for *A. frenatus* we found *wnt4*, key ligand of Wnt/β-catenin signaling that controls development and cell differentiation (79); *traf4*, an important regulator of embryogenesis and bone development (80); *hspg2*, which influences skeletal and cardiovascular development (81); and *errfi1*, that affects cell growth by regulating EGFR signaling (82). In *A. carolinensis* we detected genes associated with lipid metabolism like *plin3*, *lpin1*, *ncoa1* (83–85). For *A. auratus* we found *cib2*, associated with mechanoelectrical transduction in auditory cells (86). In *A. sagrei* we found *rab3d* involved in bone resorption (87); and *optn*, a gene associated with autoimmune and neurodegenerative disorders (88).

We combined these genes with high divergence in the regulatory regions with the genes previously identified as being under positive selection via analysis of dN/dS ratio to generate our candidate gene set. We then used STRING v11 (89) to estimate gene interaction networks for genes in our combined candidate set to obtain an integrative view of evolutionary processes that spanned both regulatory and protein divergence (Fig. 4C). Some genes with dN/dS > 1 were embedded within gene interaction networks of genes with high divergence in regulatory regions (Fig. 4C, Fig. S6). For example, in *A. carolinensis* several genes in the gene interaction network have functions associated with gene regulation, lipid metabolism, and mitochondria (Fig. 4C). The positively selected *lep* gene constitutes a central node in the gene interaction network and interacts with other elements of similar function that present high divergence in the regulatory regions.

**Figure 4.**
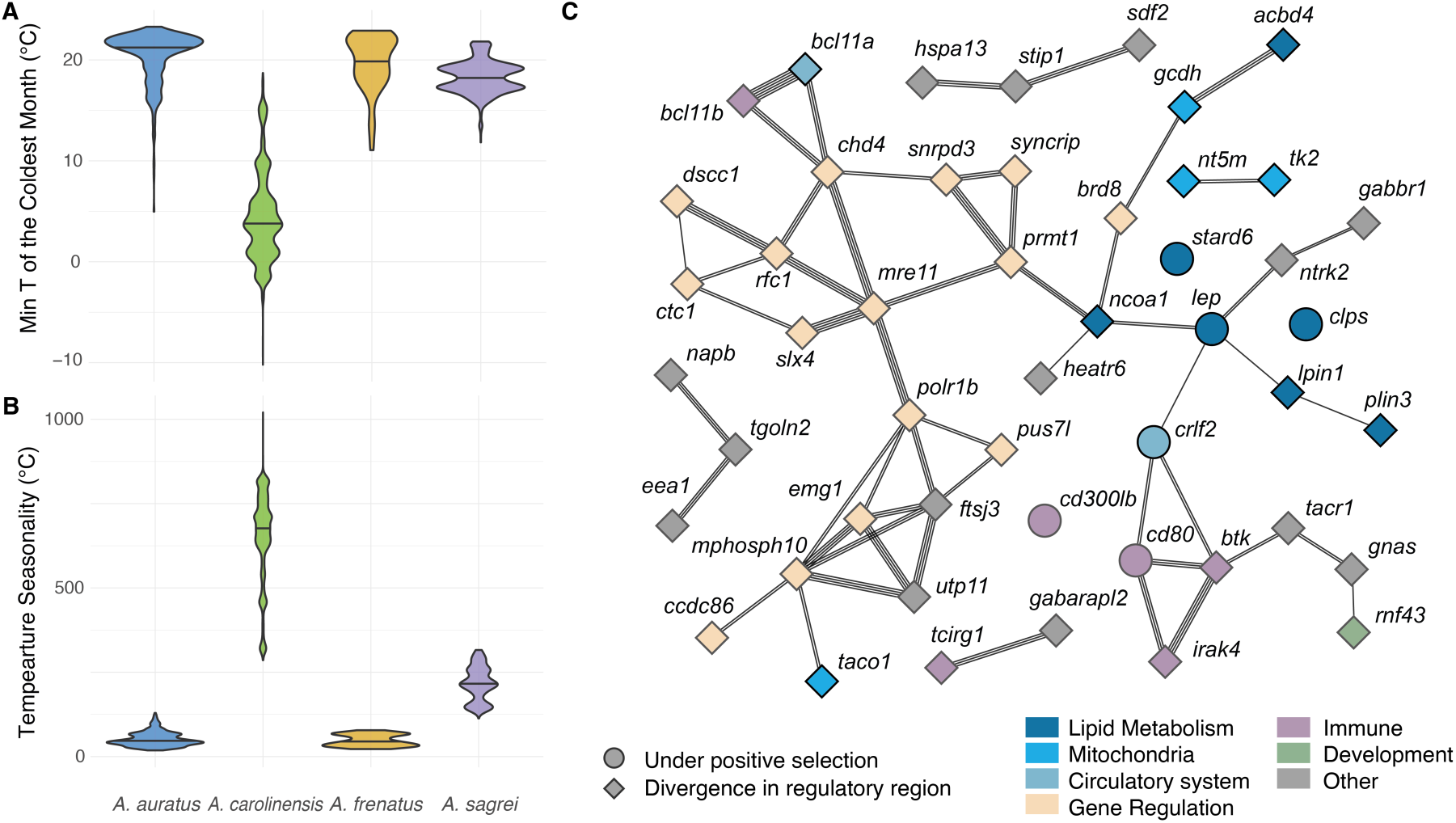
Climatic niche characterization across the native distribution range of the four studied anole species. **A.** Minimum temperature of the coldest month **B.** Temperature seasonality. *A. carolinensis* is the species inhabiting the coldest and more thermally seasonal environments. **C.** Gene interaction network for the genes under natural selection and genes with high divergence on the promoter region for *A. carolinensis*. Line thickness represents the number of multiple evidence supporting the interaction between two genes.

### Association with phenotypic traits

We characterized the realized climatic niche across the native distribution for the four *Anolis* species and detected that they have different climatic niches (Fig. S7). Among them, *A. carolinensis* occupies the coldest (Fig. 4A) and most thermally seasonal (Fig. 4B) environments. This is in accordance with the genes under selection and regulatory divergence mostly associated with biological functions that could influence cold tolerance such as lipid metabolism, mitochondria, and the circulatory system (Fig. 4C).

The morphology of the four focal species was also analyzed (Fig. S8). *Anolis frenatus* is distinct in its larger body size (Fig. 5A). The genes *mtpn* and *pdzk1ip1* were under selection in *A. frenatus* with respect to the other three species and could influence its larger body size (Fig. 5C). In contrast, *A. auratus* is characterized by its unique tail elongation (Fig. 5B). We explored the morphology of the caudal vertebrae, and we found that the long tail in *A. auratus* is associated with an elongation of the caudal vertebrae rather than an increase in the number of vertebrae when compared to the other species (Fig. 5D). The relative length of the trunk vertebrae of *A. auratus* did not differ from the other species (Fig. S9). *Anolis frenatus* also features a longer tail and longer caudal vertebrae than *A. sagrei* and *A. carolinensis*, but not as long as *A. auratus* (Fig. 5D). Among the genes under selection in *A. auratus* we detected *ramp2* and *dcdc1*, which could be associated with the vertebral elongation phenotype (Fig. 5E).

**Figure 5.**
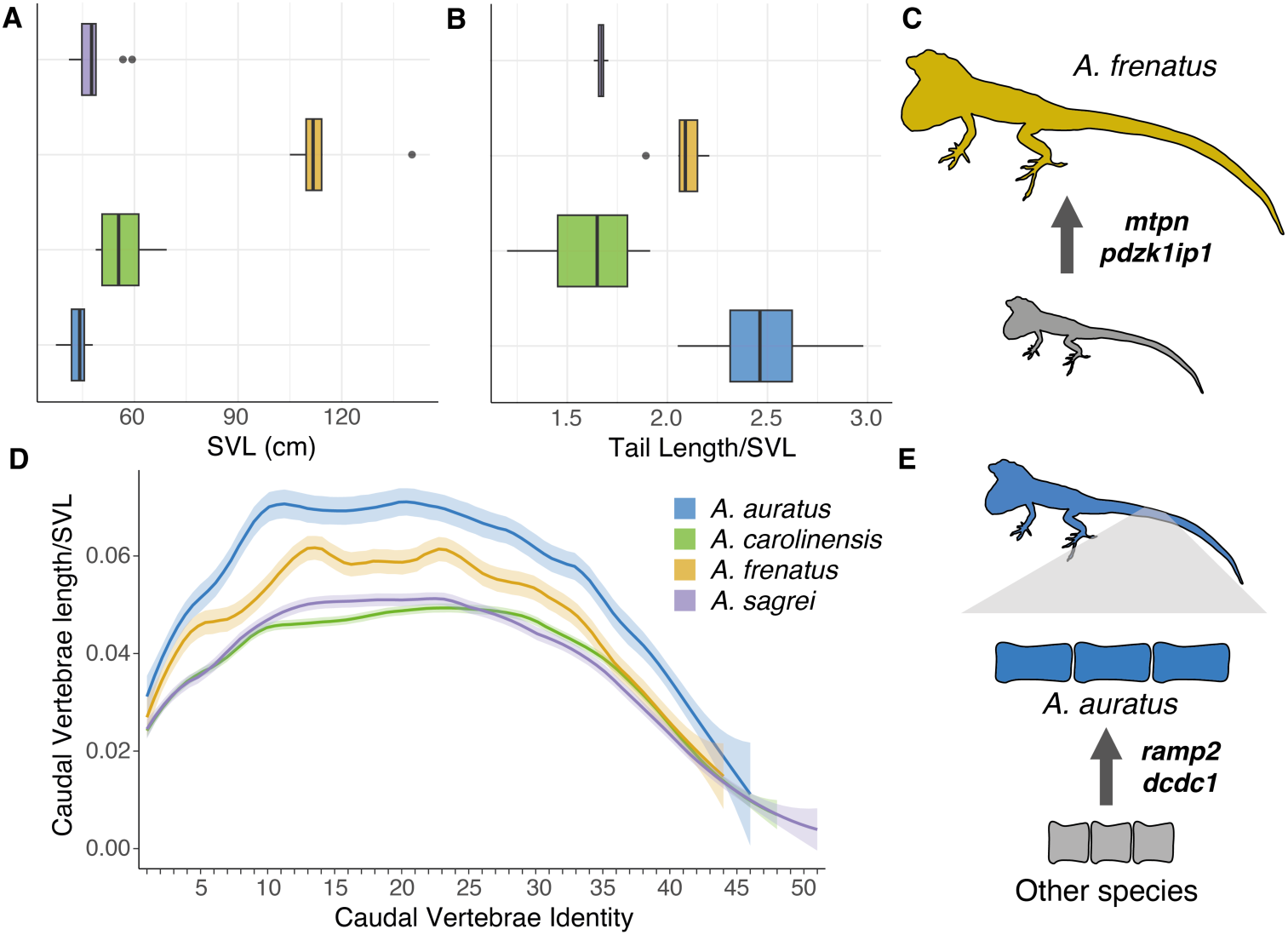
Morphological variation in distinctive traits of the analyzed species. **A.** *A. frenatus* stands out for its large body size. **B.** *A. auratus* its characterized by a long tail. **C.** Selection on the *mtpn* and *pdzk1ip1* genes could influence *A. frenatus* body size. **D.** The long tail in *A. auratus* is caused by an elongation of the caudal vertebrae rather than the addition of more vertebrae. **E.** Selection on *ramp2* and *dcdc1* could influence the vertebral elongation in *A. auratus*.

## Discussion

Genomic features can influence speciation and promote phenotypic variation within adaptive radiations (1,90). Here, we explored the genomic features, namely chromosomal rearrangements and repeat element concentration, potentially contributing to diversity and phenotypic disparity within the *Anolis* radiation. Our results indicate that major structural rearrangements, high densities of TEs around developmental genes, and signatures of natural selection and divergence on regulatory regions are associated with genes that could affect the unique phenotypes in these *Anolis* species.

### Major structural rearrangements within Anolis

Our synteny analysis detected major chromosomal rearrangements within *Anolis*. Chromosome fissions and fusions have been previously described as highly relevant in anoles (43,44), but we also identified an important number of translocations, inversions, and deletions among the analyzed species. Chromosome-level structural variations can directly influence speciation by disrupting meiosis in heterozygotes and reducing fertility in hybrids or generating barriers to gene flow (91,92). Moreover, they can modify the gene regulation and recombination patterns (10,93).

Chromosomes 1, 2 and 3 presented a substantial rearrangement in *A. frenatus* (Fig. 2B). Hi-C analysis suggests this is a true rearrangement and not a technical artifact given that contact maps show strong within-chromosome interactions and little to no interactions between these chromosomes (Fig. 2C, Fig. S2). This rearrangement was absent in the other anole species and the outgroups, which suggests this chromosomal mutation occurred in this lineage. *Anolis frenatus* is part of the basal *Dactyloa* clade of *Anolis* (Fig. 1A; (36). Chromosomes 1, 2, and 3 are bigger in other *Dactyloa* anoles when compared to non-*Dactyloa* karyotypes (Table S10), which suggests this mutation could be shared among dactyloans and evolved early during the lineage history. Chromosomal breaks in *A. frenatus* were located in areas with a high density of genes with developmental functions (Fig. 2D), including some genes highly relevant to skeletal and muscle development and growth like *axin2*, *bmp2*, *ddit3*, and *twist2* (56–59). This suggests adaptive structural rearrangements in *A. frenatus* (and potentially other *Dactyloa*) that could potentially have influenced the evolution of body size and morphology. Thus, these major structural rearrangements could have altered the gene regulation patterns of the developmental genes adjacent to them (10,93), and hence enhanced the evolution of phenotypic variability in anoles.

Our results also allowed us to explore patterns of sex chromosome evolution across *Anolis*. Anoles share a single ancestral XY sex chromosome system but have commonly experienced chromosomal fission and fusion, including fusions that involve sex chromosomes (44,94). In *A. sagrei* the X chromosome (scaffold 7) has been reported as the fusion of chromosomes 9, 12, 13 and 18 from *A. carolinensis* (45,46,60). Further, Giovannotti et al. (46) described chromosome 7 homology between *A. sagrei* and *A. valencienni*, both belonging to the *Norops* clade of anoles. Our results are consistent with this finding (Fig. 2A; Fig. S3) and expand the homology for the sex chromosome to all three analyzed *Norops*-clade anoles (*A. auratus*, *A. apletophallus* and *A. sagrei*), with only within-chromosome structural changes such as inversions and deletions differing among these species (Fig. S3). *Norops* is one of the most diverse clades within *Anolis* (36) with ∼200 species. Our findings suggest that the X-autosome fusions detected in *Anolis sagrei* arose early in the clade and highlight the relevance of sex chromosome evolution for anole diversification (44,94).

### Key developmental genes in repeat-rich regions in Anolis and other pleurodonts

We quantified the density of repeat elements through the genome for four anoles and two phrynosomatid outgroup species. Repeat elements can be a source of genetic variation because they can modify gene regulation patterns, be a source of mutations, and trigger structural rearrangements (7,95). We found a high density of repeats associated with key developmental genes such as *notch4*, *fgf11*, and the *hoxB*, *hoxC* and *hoxD* clusters in *Anolis* and the outgroups (Fig. 3; (31,62,63)). Furthermore, the enrichment analysis detected that genes located in repeat-rich regions were mostly associated with developmental and regulatory functions (Fig. S4, Table S6). Feiner (47,48) reported that anoles have an accumulation of repeats around the *hoxB* and *hoxC* gene clusters when compared against other more distantly related squamates but did not include other pleurodont lizards such as the prhynosomatids *U. nigricaudus* and *P. platyrhinos*. Thus, our analysis expands this pattern of repeat element accumulation to other genes that also affect development (Table S5) and indicates that this is not a feature exclusive to *Anolis* but is also present in other species from the Pleurodonta clade of Iguania. Pleurodont lizards include some of the most diverse vertebrate genera with respect to species number and morphological variation (e.g. *Anolis*, *Liolaemus*, *Sceloporus*; (96,97)). Therefore, the accumulation of repeat elements around developmental genes could be a source of genetic variation that fueled morphological innovation in these pleurodont groups (48). Exploring the potential effects of the repeat accumulation on genetic and phenotypic variation for these lizard groups is key to understanding whether TE dynamics contribute to their evolvability and diversification. However, additional genomes assembled from within pleurodonts and other iguanians are needed to identify specifically when this pattern arose.

### Selection on coding regions and regulatory divergence associate with unique phenotypes in Anolis

For the analyzed species we detected some candidate genes under selection and genes with high divergence in their regulatory regions, potentially associated with their unique phenotypes (Fig. 4, Fig. S6). *Anolis carolinensis* presented signatures of selection and high regulatory divergence on genes potentially driving cold adaptation. In general, ectotherm adaptation to cold environments involves physiological processes of oxygen consumption and blood circulation (28,98). Among the genes under selection in *A. carolinensis*, leptin (*lep*) was a central node in the gene interaction network (Fig. 4). Moreover, other genes associated with lipid metabolism (e.g. *clps*, *stard6*, *ncoa1*, *lpin1*, *plin3*) were also identified in our analysis. Lipid metabolism has been proposed as a potential thermal adaptation in ectotherms (99). For instance, it could be an alternative for energy obtention during cold seasons with lower resource availability (100), or it could be associated with changes in cell membrane composition impacting fluidity in colder temperatures (101). Genes associated with lipid metabolism have been identified as undergoing accelerated evolution when comparing Cuban anole species with different thermal biology (40). Furthermore, genes interacting with leptin and involved in lipid metabolism have been identified as under-selection in *A. cybotes* populations inhabiting cold high-elevation environments (102). Therefore, it is possible that changes in lipid metabolism could constitute an adaptation to cold environments in *A. carolinensis*. We also detected divergence in the regulatory region of genes associated with the circulatory system and mitochondria (Fig. 4). Populations of *A. carolinensis* inhabiting colder environments show lower oxygen consumption rates, and signatures of selection and changes in the expression of genes associated with the circulatory system (28,103). Thus, changes in these genes could enhance oxygen intake for low oxygen availability under cold temperatures in *A. carolinensis* versus other anole species.

*A. auratus* stands out for its long tail. This species is usually found on the grass in dense vegetation patches, and a long tail may provide better balance when walking or jumping across narrow perches (26,27). Body elongation is a convergent phenotype in several reptiles, and most species develop longer bodies through the addition of vertebrae (29). However, the extremely long tail in *A. auratus* is achieved by elongation of the caudal vertebrae rather than the addition of more segments (Fig. 5D). The longest caudal vertebrae in *A. auratus* are located distal to the ninth caudal vertebrae (e.g., Ca10-21, Fig. 5D). In anoles, the *m. caudofemoralis longus* originates from the proximal caudal vertebrae (e.g. Ca2-8 in *A. sagrei*, (104); Ca2-9 in *A. heterodermus*, *A. tolimensis, and A. valencienni*, (104,105); and Ca3-8 in *A. carolinensis*, (106). This primary hip joint extensor is essential for locomotion and may also assist with lateral flexion of the tail when the hindlimb is fixed (106). Therefore, caudal vertebral elongation in *A. auratus* is most pronounced in a region of the tail that is less functionally constrained. The pattern of caudal vertebrae elongation in *A. auratus* is similar to that seen in the tail of arboreal *Peromyscus maniculatus* (107), the cervical vertebrae of giraffes (108), the trunk of some plethodontid salamanders (109) and some fish species (110). Among the mechanisms that could determine caudal vertebral elongation are genes associated with axial development and determinants of the caudal region such as the *hox13* genes, *fgf8*, or *fgfr1* (30,31,107,108,111). In our genetic data, we detected signatures of selection in *ramp2* and *dcdc1*, which influence bone development (67,69). Heterozygote knockout mice for *ramp2* present skeletal abnormalities such as lower bone density and delayed development of the lumbar vertebrae, producing a similar pattern of vertebral elongation (112). In *Peromyscus maniculatus*, *dcdc1* is located within a locus associated with tail length (107). Thus, the mutations in these genes could contribute to the unique tail phenotype in *A. auratus*.

Finally, *A. frenatus* is characterized by a large body size and relatively long limbs. In general, vertebrate body size is determined by genes associated with insulin growth factors and growth hormone pathways (32,34,113,114). Our selection analysis identified some candidate genes potentially associated with large body size in *A. frenatus*. We detected signatures of selection on the *mtpn* and *pdzk1ip1* genes, both involved in muscle development, growth, and morphogenesis (64–66,115,116). Injection of *mtpn* in mice produces increased body and muscle weights (117). Moreover, among the genes that presented high divergence in regulatory regions for *A. frenatus* we identified other genes highly relevant for development. For instance, *wnt4* can be modulated by the growth hormone (118), and mice with overexpression of *wnt4* present dwarfism (119). Further, knockout mice for *traf4* show reduced body weight than wildtype mice (120).

Overall, the genes under selection and with high divergence in their regulatory regions perform relevant biological functions that could affect the phenotypes of the analyzed species. This indicates that the combination of mechanisms acting at different hierarchical levels can aid in the generation of adaptive phenotypes in anoles. Changes in regulatory regions could provide more evolvability than changes in protein-coding sequences that are in general more constrained to mutations given their biological function (40,121). Therefore, exploring the effects of regulatory sequence divergence and regulatory RNAs on gene expression and their impacts on species traits is key to understanding how this feature could promote anole phenotypic diversity.

## Conclusions

Our analysis of highly contiguous genome assemblies of four anole species allowed us to identify genomic features that could contribute to the extensive phenotypic variation among *Anolis* species. In *Anolis*, chromosome-level structural rearrangements could directly generate reproductive isolation, and affect the gene regulation patterns of genes relevant to development and morphological configuration. Further, a high density of repeat elements close to key developmental genes could also contribute to variation in the expression of such genes. Finally, natural selection on few coding sequences but relevant to species traits, in addition to divergence in regulatory regions could also play a role in shaping phenotypic diversity. The interaction between these genomic characteristics and selection pressures potentially enabled the evolution of disparate phenotypes within anoles, but further analysis of a wider sample of high-quality genomes would help to formally test this hypothesis. We highlight that besides ecological opportunity, genomic features can also be extremely relevant for promoting adaptive radiation.

## Methods

### Sampling and type specimens

The *A. auratus* specimen was collected in Gamboa, Panama, and the *A. frenatus* specimen in Soberania National Park, Panama (Collecting Permits: SE/A-33-11 and SC/A-21-12, Autoridad Nacional de Ambiente, ANAM, Republic of Panama; IACUC Protocol: 2011-0616-2014-07 Smithsonian Tropical Research Institute). Additional samples of *A. carolinensis* and *A. sagrei* obtained from the Sullivan Company (Nashville, TN) and Marcus Cantos Reptiles (Fort Myers, FL) were included for morphological analyses (IACUC Protocol: Arizona State University 19-1053R and 12-1247R). Table S1 shows the number of individuals collected per species and locations used for reference genome assemblies and morphological analyses. Specimens were euthanized by intracoelomic injection of sodium pentobarbital (IACUC Protocols 09-1053R, 12-1274R, and 15-1416R ASU). The type specimens for the *A. auratus* and *A. frenatus* reference genomes corresponded to adult females.

### Reference Genomes

We generated new reference genomes for *A. auratus* and *A. frenatus*. Skeletal muscle from the *A. auratus* type specimen, and liver and heart from *A. frenatus* type specimen were sent for DNA extraction and whole genome sequencing. The RUC_Aaur_2 and RUC_Afre_2 genomes were sequenced by Dovetail Genomics on an Illumina PE150 platform, de novo assembled with meraculous v2.2.2.5 (122). HiRise v2.1.6-072ca03871cc (123) scaffolding was performed with Chicago and Hi-C chromatic conformation capture libraries. The published genome assemblies and annotations of *A. carolinensis* (AnoCar2.0, (50); and Hi-C assembly from DNAzoo, (51,52)) and *A. sagrei* (AnoSag2.1, (45)) were included for comparative genomic analyses. Table 1 shows the assembly statistics for the four *Anolis* genomes. Additionally, we included the reference genome of the phrynosomatids *Phrynosoma platyrhinos* (MUOH_PhPlat_1.1, (54)) and *Urosaurus nigricaudus* (ASU_Uro_nig_1, (53)) for some comparative genomic analyses.

A genome annotation was generated for *A. auratus* and *A. frenatus*. For each species, repeats were identified on the genome sequences by using RepeatModeler v2.0.1 (124), and then repeat elements were soft-masked on the assembly with RepeatMasker v4.1.1 (125). To aid in annotation we generated a *de novo* transcriptome for each species using tail and ovary/yolk for *A. auratus* and brain and ovary for *A. frenatus*. Tissue samples were collected from the same animals used for genome sequencing. Tissues were sent to the Yale Center for Genomic Analyses (YCGA; West Haven, CT) for RNA extraction, cDNA poly-A-enriched Illumina library preparation and sequencing on an Illumina NovaSeq S4 platform using 150-bp paired end reads. Read quality was assessed with FastQC v0.11.7 (126), and reads were trimmed with Trimgalore v0.6.8 (127). Then a *de novo* transcriptome assembly was generated with Trinity v2.12.0 (128). The generated transcriptomes were used as evidence for each species genome annotation respectively.

Multiple iterations of Maker v3.01.03 (49) were run to annotate the genomes. We used the species-specific transcripts, and the protein-coding sequences from *A. carolinensis* and *A. sagrei* as evidence. A first round of Maker was run for aligning and mapping transcript and protein evidence. Then, two additional rounds of *ab initio* gene model prediction using Augustus v3.4.0 (129) and SNAP v2006-07-28 (130) were run. After each round of Maker, the Annotation Edit Distance (AED) was recorded, and annotation completeness was assessed with BUSCO v5.4.2 (131) on the predicted transcripts obtained from Maker, comparing against the eukaryote and sauropsid gene datasets.

### Chromosome-level structural rearrangements

We investigated synteny among the main scaffolds from the four analyzed *Anolis* species along with the phrynosomatids *P. platyrhinos* and *U. nigricaudus* by *in silico* chromosome painting. For this analysis, we used the high contiguity DNAzoo Hi-C-scaffolded genome assembly of *A. carolinensis* (51,52). All species were compared against the *A. sagrei* genome as a reference, because it is the species with the most contiguous and complete genome among our samples (45). The first 14 scaffolds from *A. sagrei*, representative of its chromosomes, were split in chunks of 100 bp with “faSplit” v438 from the UCSC Bioinformatic Utilities (132). Then, we used blastn v2.10.0 (133) to map each fragment onto the 5 other species’ genome. We retained matches with at least 50 bp length, and that were contiguous in at least 5 matches (54). To further explore the chromosome X evolution within *Anolis*, we compared chromosome 7 from *A. sagrei* to the closely related *A. apletophallus* (61) following the same methodology.

### Hi-C data analysis

Link density histograms were generated with Juicer v2.0 (134) by mapping paired reads from the Hi-C libraries for *A. auratus* and *A. frenatus* to the finished genome assembly to assess chromatin conformation and to validate our chromosomal rearrangements. Hi-C contact maps were visualized with Juicebox v1.9.8 (52).

### Developmental genes located in A. frenatus scaffolds 1, 2 and 3 rearrangement

We explored which genes were located adjected to the rearrangement among scaffolds 1, 2, and 3 detected between *A. frenatus* and *A. sagrei*. For this, we pulled from the *A. sagrei* annotation the genes located within 1 Mbp from the scaffold breakpoints identified with the synteny analysis. We performed an enrichment analysis on the genes located in these regions with g:Profiler ve111_eg58_p18_30541362 (55) to assess which biological processes were overrepresented in that gene list. Then, we extracted the list of genes present in scaffolds 1, 2, and 3 in *A. frenatus* to identify if the chromosomal breaks were located in hotspots of genes with developmental function. We identified and extracted all the GO terms included in the list of genes located on each scaffold with g:Profiler, and we retained only the genes matching GO terms that included any of the keywords: “development”, “morpho”, “growth” or “organ”. We then calculated the number of genes with those developmental functions along each chromosome in 500 kbp windows in R v4.1.2 (135) with a custom script.

### Repeat density through the genomes

For *A. auratus*, *A. frenatus*, *A. sagrei*, *U. nigricaudus* and *P. platyrhinos* we calculated the repeat density for each one of the largest 6 scaffolds. The number of repeats was calculated in 500 kbp windows, and we retained repeats longer than 50 bp and with a score value over 10 (47). Then, we selected the 500 kbp windows corresponding to the highest 1% of repeat density per scaffold for each species with a custom script in R, and identified which genes were located in those high repeat density regions using the respective genome annotations. An enrichment analysis was performed to identify the most represented gene ontology (GO) categories on the list of genes situated in high repeat regions for each species with g:Profiler, and the enriched GO terms were semantically organized and visualized with Revigo v1.8.1 (136).

### Identification of genes under positive selection and regulatory elements with high divergence

We looked for potential genes under positive selection among the four *Anolis* species by calculating the ratio between non-synonymous to synonymous mutations (dN/dS) between orthologs from species pairs with the “orthologr” package (137) in R using the Comeron’s (138) method. To identify genes positively selected in each species, we retained the genes overlapping in at least 2 out of 3 comparisons between the focal and the other three species. Then, an enrichment analysis was performed on that gene list with g:Profiler to identify the most represented GO terms for each species.

To assess regulatory regions with high divergence we focused on the 1000 kb upstream of the transcription start, which includes the promoter region (78). We compared orthologs between species pairs previously identified with “orthologr” in R. Each ortholog pair was aligned with mafft v7.520 (139) and the genetic distance between aligned orthologs was estimated with the “bio3d” package (140) in R. We considered the genes with the top 1% of genetic distance as genes with the highest divergence in their regulatory regions between species pairs. For each species we retained the genes overlapping in at least 2 out of 3 comparisons. Finally, we used STRING v12.0 (89) to evaluate gene interactions among the genes under selection and the genes with high regulatory divergence for each species.

### Climatic niche analyses

For each species, occurrence records were obtained from the Global Biodiversity Information Facility (141). Occurrences were deduplicated and manually curated to accurately represent the native distribution of each species. Raster data for the 19 bioclimatic variables was obtained from the WorldClim v2 database (37). For each occurrence point, the corresponding values of the 19 bioclimatic variables were obtained. We compared the climatic niche between the four analyzed species. Climatic variation was visualized with a Principal Component Analysis (PCA) in R, and the main variables differentiating species were identified.

### Morphological analyses

Additional samples for the four *Anolis* species were included for morphological analyses. Skeletal data was obtained from osteological preparations following Tollis et al. (2018) modification of amphibian protocols, or from micro-computed tomography (microCT) images collected in a Siemens Inveon micro-CT scanner at the RII Translational Bioimaging Resource at the University of Arizona (Table S1). For skeletal preparations, individuals were photographed with a scale in a stereodissecting microscope (Nikon SMZ800 with Coolpix 995 digital camera), and morphological traits were measured with ImageJ v1.53k (142). For microCT scans, digital images were analyzed and measured with InVesalius v3.1.1 (143). For each species we measured snout–vent length (SVL), axilla– groin distance (AGD), forelimb total length (FLL), forelimb autopod length, forelimb stylopod length, forelimb zeugopod length, hindlimb total length (HLL), hindlimb autopod length, hindlimb stylopod length, hindlimb zeugopod length, head width (HW), head length (HL), head height (HH) and tail length (TL). We also analyzed the osteology of the caudal vertebrae for the four *Anolis* species. For this, we measured the distance from the distal end of the cotyle to the proximal tip of the condyle on each caudal vertebra. All measurements apart from SVL were standardized by dividing by the distance between the snout to the end of the sacral vertebrae as an approximation to body size. We compared microCT and skeletal preparation measurements with a paired T-test to assess possible bias in the sampling methodology (Table S11).

## Data Availability

All raw read files have been accessioned to the NCBI SRA under BioProject #PRJNA1096315. Final genome assemblies and annotations have been accessioned to the Harvard Dataverse (https://doi.org/10.7910/DVN/F9NDWL).

